# Characteristics of respiratory measures in young adults scanned at rest, including systematic changes and “missed” respiratory events

**DOI:** 10.1101/613851

**Authors:** Jonathan D. Power, Benjamin M. Silver, Alex Martin, Rebecca M. Jones

**Affiliations:** Sackler Institute for Developmental Psychobiology, Department of Psychiatry, Weill Cornell Medicine, 1300 York Avenue, Box 140, New York, NY 10065 USA; National Institute for Mental Health, 10 Center Dr. Bethesda, MD 20814 USA

## Abstract

Breathing rate and depth influence the concentration of carbon dioxide in the blood, altering cerebral blood flow and thus functional magnetic resonance imaging (fMRI) signals. Such respiratory fluctuations can have substantial influence in studies of fMRI signal covariance in subjects at rest, the so-called “resting state functional connectivity” technique. If respiration is monitored during fMRI scanning, it is typically done using a belt about the subject’s abdomen to record abdominal circumference. Several measures have been derived from these belt records, including the windowed envelope of the waveform (ENV), the windowed variance in the waveform (respiration variation, RV), and a measure of the amplitude of each breath divided by the cycle time of the breath (respiration volume per time, RVT). Any attempt to gauge respiratory contributions to fMRI signals requires a respiratory measure, but little is known about how these measures compare to each other, or how they perform beyond the small studies in which they were initially proposed. In this paper, we examine the properties of these measures in hundreds of healthy young adults scanned for an hour each at rest, a subset of the Human Connectome Project chosen for having high-quality physiological records. We find: 1) ENV, RV, and RVT are all similar, though ENV and RV are more similar to each other than to RVT; 2) respiratory events like deep breaths exhibit characteristic fMRI signal changes, head motions, and image quality abnormalities time-locked to deep breaths evident in the belt traces; 3) all measures can “miss” respiratory events evident in the belt traces; 4) RVT “misses” deep breaths (i.e., yawns and sighs) more than ENV or RV; 5) all respiratory measures change systematically over the course of a 14.4-minute scan, decreasing in mean value. We discuss the implication of these findings for the literature, and ways to move forward in modeling respiratory influences on fMRI scans.

**Highlights:**

- Examines 3 respiratory measures in resting state fMRI scans of healthy young adults
- All respiratory measures “miss” respiratory events, some more than others
- Respiration volume per time (RVT) frequently “misses” deep breaths
- All respiratory measures decrease systematically over 14.4 minute scans
- Systematic decreases are due to decreased breathing depth and rate

## Introduction

Modulation of functional magnetic resonance imaging (fMRI) signals by respiration has come under increased scrutiny in recent years, largely due to the emergence of “resting state functional connectivity” approaches to signal analysis, which tend to examine signals in terms of covariance rather than in terms of pre-defined events modeled in time. Changes in breathing depth and rate modulate the amount of carbon dioxide (CO_2_) exhaled, and thereby alter the concentration of CO_2_ in the bloodstream (pCO_2_). Cerebral blood flow is controlled by homeostatic loops governed mainly by pCO_2_, such that increased pCO_2_ causes increased cerebral blood flow (Hall, 2016). A typical baseline pCO_2_ is 35-45 mm Hg. Increases of ~5 mm Hg caused by decreased breathing rate or depth, or inhalation of CO_2_-enriched gas mixtures, cause cerebral blood flows to increase on the order of 50%, which in turn cause large multiple-percent blood oxygen level dependent (BOLD) signals changes throughout the brain (Bright et al., 2009; Ito et al., 2003; Kastrup et al., 1999; Poulin et al., 1996). In this manner, breathing patterns can alter fMRI signals.

The timescale of pCO_2_-induced BOLD signal changes is long compared to that of typical task responses. “Evoked” fMRI responses to transient single deep breaths tend to span 30-40 seconds (Birn et al., 2006; Birn et al., 2008; Chang and Glover, 2009; Power et al., 2018; Power et al., 2017b) (see Figure 1), whereas the evoked fMRI responses to experimental button presses or transient visual stimuli are typically over within 15 seconds of an event (which is why brief task events are usually modeled over comparably brief periods (Buxton et al., 2004)). Because breathing causes large signal modulations spanning relatively long times, breathing can have considerable influence on fMRI signal covariance. In task studies, the temporal design permits one to average away physiological “noise” so long as it is not correlated with the task design. However, in functional connectivity studies, there is no comparable way to eliminate such noise without explicit effort.

**Figure 1:**
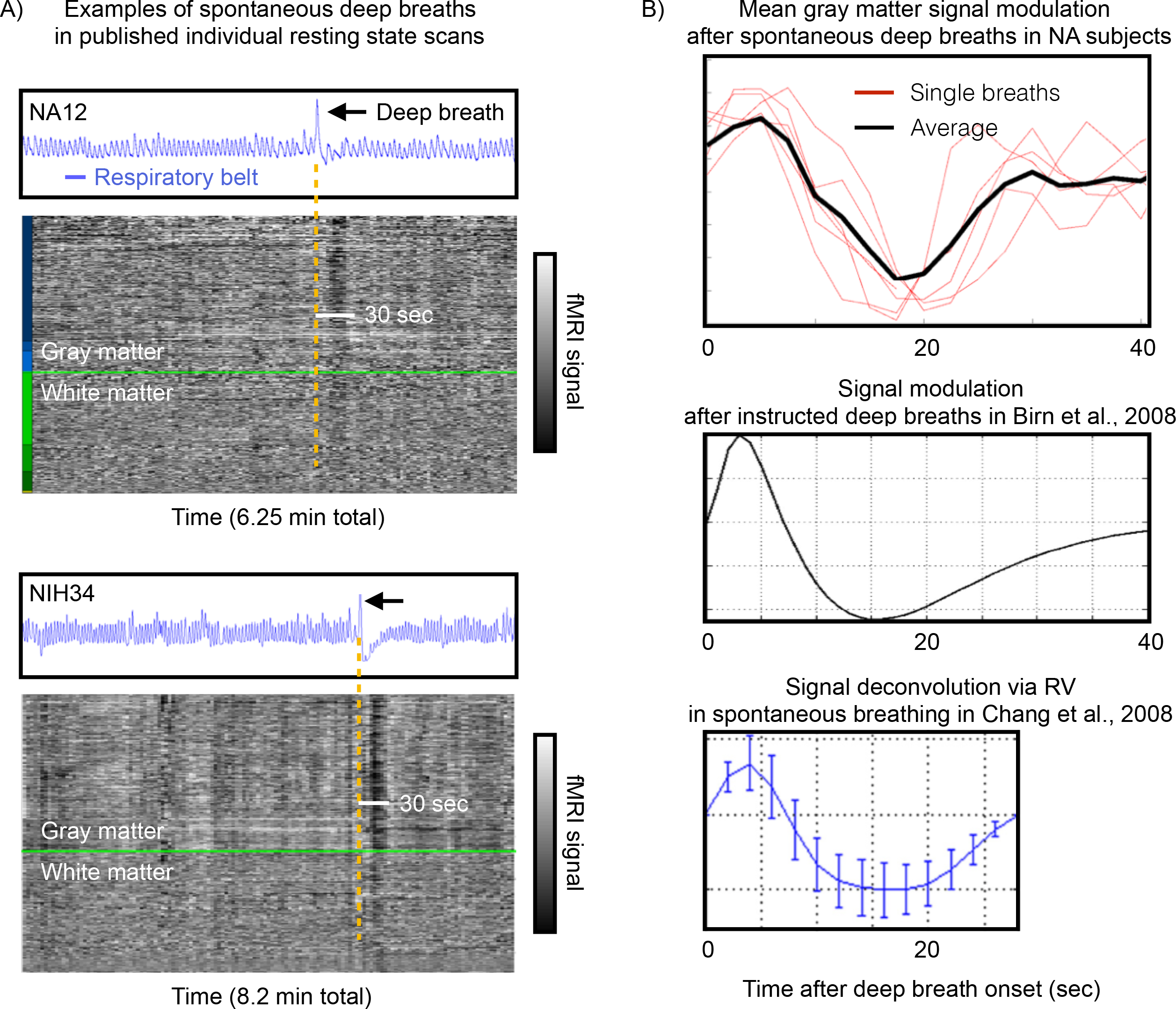
Examples of respiratory traces and effects of deep breaths in published datasets. A) For two individual scans published in {Power, 2018} and {Power, 2017}, blue traces show respiratory belt records, and the gray-scale heat maps show all in-brain fMRI signals ordered by anatomic compartment, with gray matter above the bright green line and white matter below the bright green line. A single deep breath is evident in each scan, and accompanying fMRI signal modulations last about 30 seconds in both scans. B) Top panel shows mean gray matter signals in the 40 seconds after isolated deep breaths in subjects at rest in {Power, 2018}, middle panel shows the average response to instructed single breaths in {Birn, 2008}, and bottom panel shows respiratory response function derived by deconvolution of multiple physiological parameters (including RV) from fMRI signals in subjects at rest {Chang, 2009}.

Any effort to identify respiratory influences on fMRI signals – whether in task or task-free settings – must begin with a measure of respiration. In modern studies of respiratory physiology, it is common practice to take multiple kinds of measures at once, providing both unique and redundant information, and enabling high-confidence identification and characterization of respiratory phenomena. Typical noninvasive measures of respiration are optical or inductance respiratory plethysmography, which involve simultaneous optical or mechanical measurement of thoracic and abdominal excursions (since either compartment can move air nearly independently). Such measures are often paired with nasal cannula or face-mask measures of air flow and end-tidal pCO_2_ (i.e., capnography) (Heinzer et al., 2015; Yumino and Bradley, 2008). These approaches can produce precise, quantitative measurement of air flow and blood gas changes, in addition to characterizing other respiratory parameters.

By contrast, in typical fMRI studies, if respiratory monitoring occurs at all, it is accomplished via a “respiratory bellows”, which is a single mechanical belt strapped about the subject’s abdomen that measures tension on the belt (see blue traces in Figure 1A). This belt has no back-up source of redundant information (e.g., a chest belt), and will index both respiratory motion and non-respiratory motions if they involve shifts of the abdomen and/or tightening of abdominal muscles. The abdominal belt does not permit quantitative measurement of air flow in the manner of calibrated plethysmography, but several semi-quantitative measurements have been proposed to index respiratory phenomena from the belt traces. These include respiration variation (RV), which is the standard deviation of the belt trace within a (6-second) window (Chang et al., 2009), the envelope of the respiratory trace (over a 10-second window, ENV) (Power et al., 2018), and the change in belt magnitude over a breath cycle (respiratory volume per time, RVT) (Birn et al., 2006).

These semi-quantitative respiratory measures (RV, ENV, and RVT) have been used to model physiological noise in fMRI signals, and their explanatory power has been greatest when such measures are convolved with “respiratory response functions” of the kind shown in Figure 1B (Birn et al., 2006; Birn et al., 2008; Chang et al., 2009; Chang and Glover, 2009; Power et al., 2017b). These respiratory response functions characterize the relatively slow signal changes that occur in the 30-40 seconds after a brief increase in ventilation. The functions are similar no matter whether they are derived from instructed deep breaths (Birn et al., 2008), from a deconvolution of multiple cardiac and pulmonary signals in task-free fMRI signals (Chang et al., 2009; Chang and Glover, 2009), or when estimated from isolated spontaneous deep breaths in subjects at rest (Power et al., 2018; Power et al., 2017b). In short, measures like ENV, RV, and RVT are intended to index respiration and respiratory “events”, and the respiratory response functions are intended to describe the consequent signal changes.

In this paper we explore the major features of respiration that these measures capture, and the ability of these respiratory measures to capture respiratory “events”. We have used all of these measures in multiple datasets and hundreds of scans over many years, and have noticed particular features of certain measures that constrain their uses in certain ways. In particular, RVT, due to its definition, often “misses” events that are evident visually in respiratory traces and that also demonstrate characteristic fMRI signal consequences of respiratory changes. More broadly, all of the measures can “miss” an event depending on the characteristics of the respiratory belt trace. We illustrate these phenomena in Human Connectome Project (HCP) data, which are attractive for the sheer quantity of data (hundreds of subjects with an hour each of resting state fMRI with accompanying respiratory belt measurements). We also describe systematic changes in the respiratory measures over a scanning period.

For readers unfamiliar with respiratory terminology, the following terms are used: *tidal breathing* is normal, cyclic breathing, *tidal volume* is the volume of air inspired in a normal breath, *minute ventilation* is the volume of air moved in a minute (respiratory rate times tidal volume), *eupnea* describes the state of normal, unlabored tidal breathing, *hyperpnea* and *hypopnea* describes increased and decreased ventilation, respectively, and *apnea* is an absence of ventilation.

## Methods

### Data

The data studied are exactly the data studied in (Power et al., 2019a).

The “900-subject” release of the Human Connectome Project data was obtained, with a focus on the following files in each subject:

Four resting state fMRI scans transformed to atlas space (in each subject’s /MNINonLinear/Results folder): {RUN}=REST1_LR, REST1_RL, REST2_LR, REST2_RL (this order is runs 1-4 in the text). rfMRI_{RUN}.nii.gz and rfMRI_{RUN}_hp2000_clean.nii.gz scans were obtained, representing minimally preprocessed and FIX-ICA-denoised data. For each of these scans, the {RUN}_Physio_log.txt and Movement_Regressors_dt.txt files were also obtained. Structural scans transformed to atlas space (in each subject’s /MNINonLinear/ folder): the T1w.nii.gz and the aparc+aseg.nii.gz files, representing the anatomical T1-weighted scan and its FreeSurfer segmentation.

### Image Processing

The aparc+aseg.nii.gz file for each subject underwent a set of serial erosions within white matter and ventricle segments, exactly as in (Power et al., 2017b). Masks of cortical gray matter, the cerebellum, and subcortical nuclei were extracted, as were serially eroded layers of superficial, deep, and deepest (with respect to distance from gray matter) masks of the white matter and ventricles. These masks, together, include all in-brain voxels of these tissue types, and are used to extract certain signals and to order signals for “gray plots” (Power, 2017).

For the purpose of making useful gray plots, because of the considerable thermal noise in HCP scans, a within-mask 6 mm FWHM Gaussian kernel was applied to the data using the above masks (illustrated for HCP data in (Power, 2017)). This blurring does not mix tissue compartments due to the use of masks.

### Parameter processing

#### Respiratory and cardiac measures

Respiratory belt and pulse oximeter traces (sampled at 400 Hz) first underwent visual inspection in their entirety to determine if the quality was sufficient for reliable peak detection, since traces are often partially or fully corrupted. Only subjects with traces deemed likely to successfully undergo peak detection in all runs were analyzed. Readers may view all physiology traces and decisions on quality in the supplemental movies of (Power, 2019)^1^.

After selection, for respiratory traces, an outlier replacement filter was used to eliminate spurious spike artifacts (Matlab command: filloutliers(resp_trace,‘linear’,‘movmedian’,100)) and the traces were then gently blurred to aid peak detection (Matlab command: smoothdata(resp_trace,‘sgolay’,400)) (a 1-second window for a 400 Hz signal). These treated respiratory traces are the ones shown in Figures.

Following prior literature, several respiratory measures were derived from the treated respiratory belt trace. First, the envelope of the trace over a 10-second window (at 400 Hz) was calculated after (Power et al., 2018) (Matlab command: envelope(zscore_resp_trace,4000,‘rms’)). Second, the RV measure, defined as the standard deviation of the treated respiratory trace within a 6-second window, was calculated following (Chang and Glover, 2009) (Matlab command: movstd(zscore_resp_trace, 2400,‘endpoints’,‘shrink’)). Finally, the RVT measure, defined for all peaks as ((peak-prior trough)/(time between peaks)), was calculated after (Birn et al., 2006). Peak detection on the trace yielded peaks (and troughs, using the inverted trace) for calculations (Matlab command: (findpeaks(zscore_resp_trace,‘minpeakdistance’,800,‘minpeakprominence’,.5)). The minimum peak distance presumes breaths occur more than 2 seconds apart. If a peak did not have a preceding trough prior to the previous peak, no value was scored at that peak. All traces and derived measures were visually checked to ensure that outliers and abnormalities would not drive results. These 3 measures were termed ENV, RV, and RVT in figures.

Pulse oximeter traces underwent z-scoring then peak detection (Matlab command: findpeaks(zscore_pulseox,‘minpeakdistance’,180,‘minpeakprominence’,.5)). Heart rate was calculated from the interval between peaks. The minimum peak distance presumes heart rates are under 133 beats per minute. Peak amplitude was calculated from the height of the peak relative to the previous trough. Cardiac traces are prone to transient disruptions when fingers move, and it is laborious to check and correct cardiac measures due to the large numbers of peaks and troughs. A limited number of cardiac records are therefore used in this report, but those select traces and their derived measures were visually checked and if necessary corrected to ensure accuracy.

#### Data quality measures

The data quality measure DVARS was calculated after (Power et al., 2012; Smyser et al., 2010) as the root mean squared value in the brain at each timepoint of all voxel timeseries differentiated in time by backwards differences. DVARS by convention is 0 at the first timepoint.

#### Head position and head motion measures

Head position was taken from the Movement_Regressors_dt.txt files. In Figures these position parameters are displayed either after subtracting the first timepoint value from the timeseries (so that all traces start at zero) or by spreading them on the y-axis by adding constants to each trace (so that each individual trace is seen clearly). Head motion was represented by Framewise Displacement (FD) measures, following (Power et al., 2012), wherein all position measures were differentiated in time by backwards differences, rotational measures were converted to arc displacement at 5 cm radius, and the sum of the absolute value of these measures was calculated. FD is typically calculated by backwards differences to the preceding timepoint (here 720 ms prior), but historically FD measures using sampling rates of 2-4 seconds were common; for comparison to such measures, FD was also calculated by backwards differences over 4 timepoints (4 * 720 ms = 2.88 seconds effective sampling rate) where indicated, using position estimates that had dominant respiratory frequencies removed. This reformulation of FD is discussed extensively in (Power et al., 2019a).

#### Frequency content

At times, the frequency content of certain signals is illustrated, such as for respiratory belt traces or for position estimates. Power spectral density estimates were generated via Welch’s method (e.g., for a respiratory trace, Matlab command: [pw pf] = pwelch(signal,[],[],[],400,‘power’)) and are shown on a semilogarithmic scale. All spectra were visually checked to ensure correct peak frequency identification.

At times and where indicated, position estimates were filtered to exclude certain frequencies using a Butterworth filter (Matlab commands for an example stopband of 0.2-0.5 Hz: TR = 0.72; nyq = (1/TR)/2; stopband=[0.2 0.5]; Wn = stopband/nyq; filtN = 10; [B,A] = butter(filtN,Wn,‘stop’); filtposition(:,column) = filtfilt(B,A,position(:,column));).

## Results

### Illustration of respiratory records and respiratory measures

Of the HCP “900 subject” release, only 440 subjects had full sets of fMRI data and full sets of physiologic data in which we believed signals were of sufficient quality that we could algorithmically obtain reliable peaks in cardiac and pulmonary traces. Only these subjects were analyzed further. Characteristics of the subjects were: age 28.6 ± 3.8 (range 22 – 36), 228 males and 216 females, BMI 26.5 ± 5.0 (range 16.5 – 43.9). Throughout the paper we will distinguish between “respiratory traces”, referring to the respiratory belt waveforms, and “respiratory measures”, meaning measures derived from the respiratory traces such as ENV, RV, and RVT.

Examples of respiratory traces and associated ENV, RV, and RVT measures are shown for full runs of 8 subjects in Figure 2. Several points are noteworthy. First, the typical frequency of respiration varies by subject. Second, within a single run, there can be wide variety in respiratory frequency (e.g., 8^th^ panel). Third, within a single run, there can be a wide variety in respiratory depth (e.g., 3^rd^ panel). Fourth, there may be frank pauses in breathing (e.g., 1^st^, 5^th^, and 7^th^ panels, gray boxes). Fifth, the ENV and RV waveforms tend to be quite similar and are both distinct from the RVT waveform (see Figure S1: across subjects, ENV and RV correlate at r = 0.85 ± 0.08, and ENV and RV correlate to RVT at r = 0.64 ± 0.20 and 0.66 ± 0.18; the difference between the first and the latter correlations are significant by two-sample t-test (p = 0 in each run)). Sixth, related to the fifth point, some abnormalities in respiratory traces are reflected mainly in ENV and RV (orange boxes in 1^st^ panel) whereas others are more captured by the RVT measure (red box in 2^nd^ panel). Such plots for all runs of all used subjects are shown in the supplemental movies^2^.

**Figure 2:**
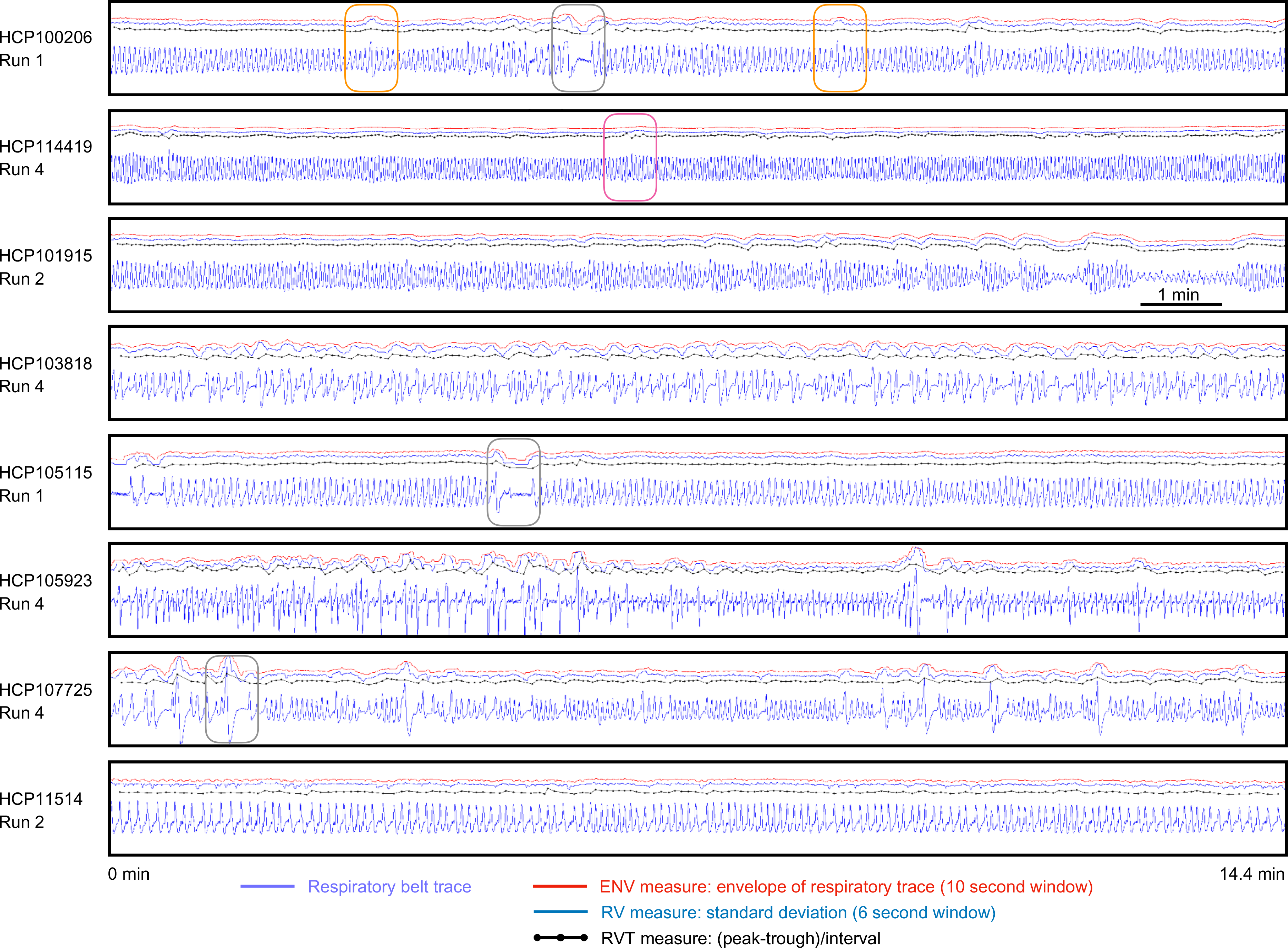
Examples of respiratory traces and derived measures. For 8 scans from 8 subjects of the HCP dataset, respiratory traces and derived ENV, RV, and RVT measures are shown. Each scan lasts 14.4 minutes. Note the wide variety of periodicity within a scan and across subjects, as well as the varied shapes of the raw respiratory belt waveforms. Often the 3 derived measures capture similar properties of the respiratory trace, such as in the third panel. But in many instances there are discrepancies: the orange boxes in the 1st panel show ENV and RV capturing deep breaths, but RVT not capturing the same phenomena. Conversely, the red box in the 2nd panel shows ENV and RV not capturing a pause in breathing, which RVT captures. Gray boxes show several pauses in breathing.

### Examination of respiration via gray plots demonstrates “missed” respiratory events

It is helpful to study respiratory traces in the context of other concurrent measures. The gray plot in Figure 3 shows a full run to orient the reader. The respiratory trace is shown in blue in the 3^rd^ panel, and exhibits rather uniform periodicity and depth (i.e., eupnea) with a few exceptions, 3 of which are marked by orange boxes. The fact that something physiologically meaningful occurred at these times is evident from the fMRI timeseries heatmap below in the 4^th^ panel, where vertical black bands representing pan-brain signal decreases follow each of the orange boxes. The heatmap shows signals from all voxels in the brain, ordered by anatomical compartment, with gray matter above the bright green line, and white matter below. The DVARS measures in the 2^nd^ panel show at these boxed times spikes in the minimally processed data (light green trace) and “DVARS dips” in the FIX-ICA-denoised data (dark green trace). These spikes and dips indicate that variance in these timepoints is changing abnormally rapidly in the minimally processed image, and abnormally slowly in the FIX-ICA-denoised images. The explanation for the dips is that some ICA components act as pseudo-delta functions to remove large amounts of variance at specific time points during denoising. Head position traces show two notable properties in the 1^st^ panel (light gray traces). First, there are zig-zags throughout the scan sharing the same periodicity as the respiratory trace, which reflects head motion and pseudomotion at the primary respiratory rate (Fair et al., 2018; Power et al., 2019b; Siegel et al., 2017). Second, at the three boxed times in question, there are sustained step changes in head position (pink arrows show starting and ending positions of a single position parameter). These position changes are captured in the FD measure of head motion, shown in bright red. Note that the FD shown here is a modified version of FD, namely one derived from comparisons of position traces across 4 TRs (2.88 seconds) that have had frequencies between 0.2-0.5 Hz filtered out. The rationale for these choices in constructing FD is detailed elsewhere (Power et al., 2019a). A 1-TR version of FD is shown in light red.

**Figure 3:**
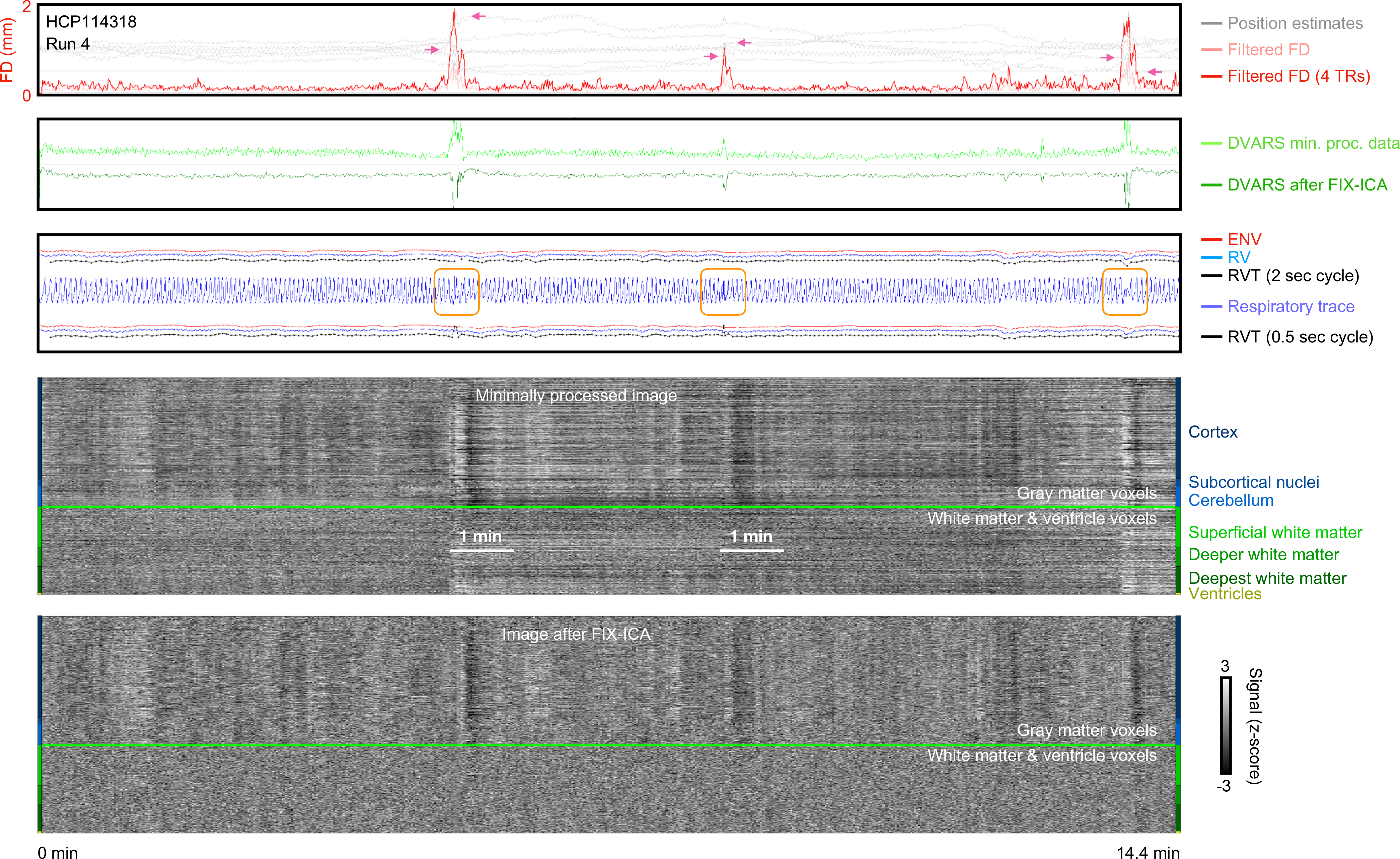
Example of scan motion, image quality measures, respiratory measures, and fMRI timeseries before and after FIX-ICA denoising. In the first panel, unfiltered position estimates are shown in light gray, illustrating large trends in head position and also periodic respiratory motion. A bright red trace shows FD calculated in 4-TR intervals of the filtered position estimates, yielding motion estimates similar to those used in the literature in recent years (effective TR = 2.88 sec); the horizontal black line denotes FD = 0.5 mm. In the second panel, DVARS (DV) calculated in minimially processed data and FIX-ICA denoised data. Note the correspondence of motion with DV spikes in minimally processed data, and the “DV dips” after FIX-ICA, indicating that the ICA procedure acted as a kind of spike regressor at these times. In the third panel, respiratory traces and derived measures are shown. Note the circled abnormalities in the belt trace, which are not always captured by the derived measures. Two versions of RVT are shown (above and below the belt trace), the upper trace using standard peak finding settings and the lower trace using a setting permitting very rapid breaths. The bottom panels show “gray plots” of all timeseries in the image, organized by anatomical compartment. Such plots for all runs of all subjects are in the online movies.

This scan illustrates important points about the respiratory measures derived from respiratory belt traces (i.e., the red ENV, blue RV, and black RVT traces immediately above the respiratory trace in the 3^rd^ panel). First, as noted earlier, all measures are generally similar, especially ENV and RV. Second, respiratory events can be “missed” by any of the measures. In the present scan, all of the measures mark the third event with a dip, but there is little indication in the measures of the first and second boxed events. In this scan the “misses” are concordant across measures, but later figures will illustrate discordant situations where some measures mark an event but others do not.

The “misses” prompt some comments on respiratory measure definition and the qualities of the respiratory traces. Tautologically, ENV and RV “missed” events because the representations of the events in the belt trace produced little or no meaningful change in the waveform envelope or its windowed variance. But, unlike the other respiratory measures, RVT depends on peak detection (RVT = peak amplitude / time between peaks), and one might object that the “misses” occurred because RVT is not optimally calculated in this subject, for there are rapid oscillations in respiratory traces at the boxed times and RVT is not calculated for each of the peaks. However, as shown below, it is probably not possible to prevent “misses” by RVT in this scan and the challenge of dealing with such traces is precisely the reason this scan was chosen for illustration.

Peak-finding settings must be chosen to operate reliably across a wide variety of respiratory waveforms, as evident in Figures 1 and 2. However, not only is there much variability in “true” respiratory waves, but the belt traces are also susceptible to abdominal contraction and motion, which can introduce rapid non-respiratory fluctuations into the trace of varying amplitude. Whereas large multi-second fluctuations of abdominal belts have face plausibility as respiratory phenomena, rapid fluctuations are less plausibly purely respiratory phenomena. For instance, amid several very small and rapid fluctuations, the first boxed event ostensibly contains two full breaths accomplished in under two seconds, an extremely rapid rate of breathing. Though there is doubtless a respiratory event at this time, it seems more likely that the subject breathed deeply but also moved and contracted the abdominal musculature to yield a rapidly fluctuating waveform, rather than suddenly breathing rapidly with odd variations in volume amidst an otherwise eupneic scan. Note that the black bands in the fMRI signals here span ~30-40 seconds, just as the deep breaths did in Figure 1. Without another source of respiratory information, however, the truth of what occurred cannot be known with certainty.

Given the uncertainty in the “meaning” of rapid oscillations in the belt traces, it is prudent to choose peak-finding settings that minimize the production of non-respiratory values, especially the outliers that would likely be generated by brief cycle times. Nearly all scans have respiratory frequencies under 0.4 Hz, which is why a 2-second breath cycle is the standard setting in this paper. If more rapid peak detection is permitted, a greater number of high RVT values will be generated, and fewer low RVT values will be generated. Figure 3 illustrates this occurrence by re-presenting the RVT measure while permitting peaks to exist 0.5 seconds apart (see red, blue and black traces under the respiratory trace), now yielding prominent spikes in the RVT trace at the first two boxed events. Note, however, that the abnormality at the third boxed event initially seen in the “standard” RVT has now largely disappeared! As stated above, preventing “misses” in this scan due to peak-finding may not be possible.

Moreover – and this will be seen repeatedly in the paper – RVT can fail to flag a respiratory event even when there is no ambiguity about proper peak-finding. RVT is designed to index volume per time, and, if breath volume scales with breath time, changes in those parameters can cancel one another out. Clear examples are shown in Figure 4 and 5 of deep breaths being unremarkable in RVT simply because deeper-than-usual breaths occur over longer-than-usual time spans. Peak-finding plays no role in such “misses”.

**Figure 4:**
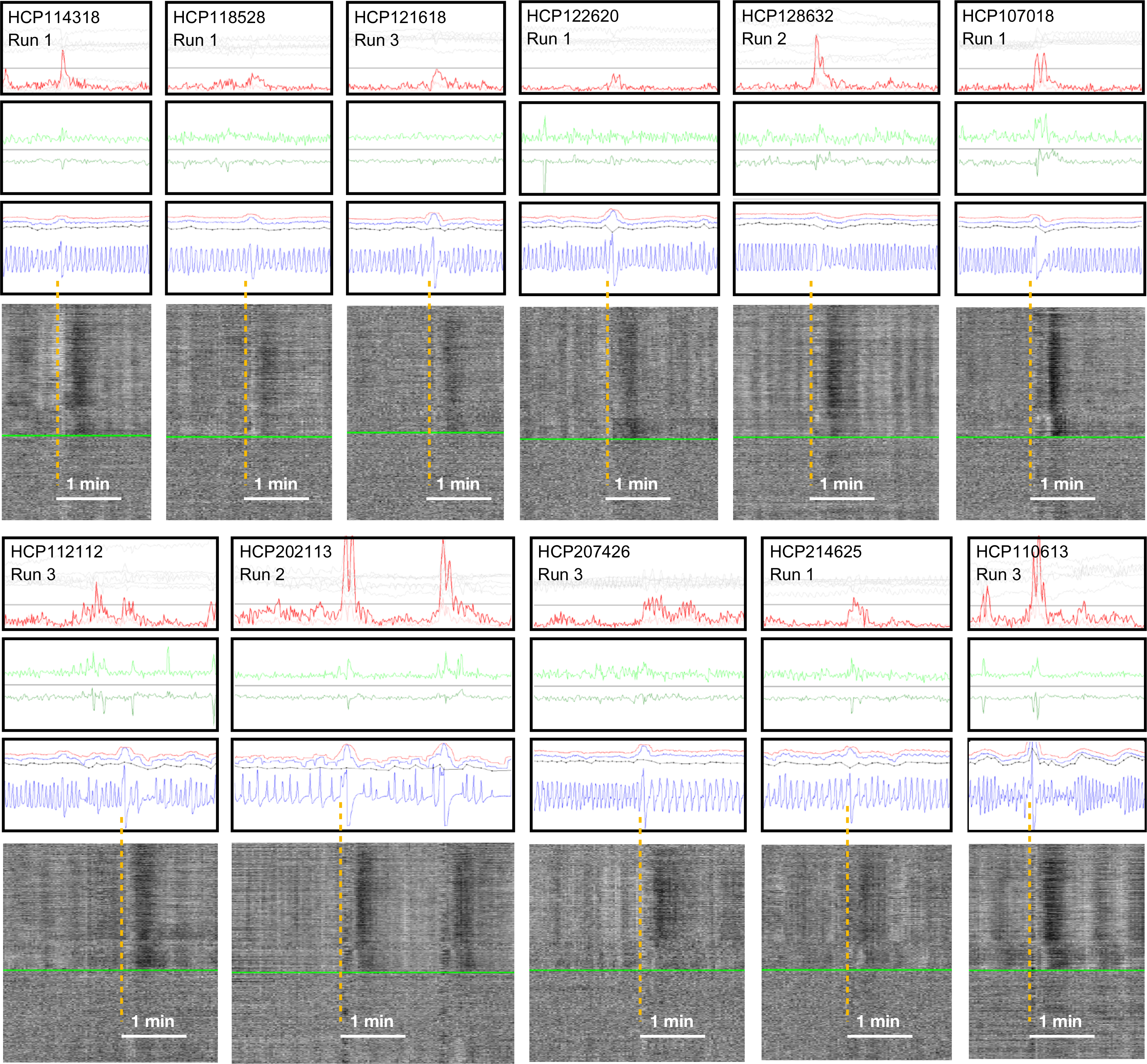
Discordance of respiratory measures during “single deep breaths”. For 11 scans with relatively isolated deep breaths (see blue respiratory traces), excerpts of figures in the style of Figure 3 are shown. The gray plots are of minimally processed data. Note that ENV (red trace) and RV (blue trace) often have bumps at the deep breaths, whereas RVT (black trace) may be flat, have a bump or spike, or may have a dip, depending on the shape of the breath. Breaths were selected by the respiratory trace and the presence of concomitant fMRI signal changes typical of deep breaths, similar to those shown in Figure 1, not on the basis of motion or DVARS or respiratory measures.

**Figure 5:**
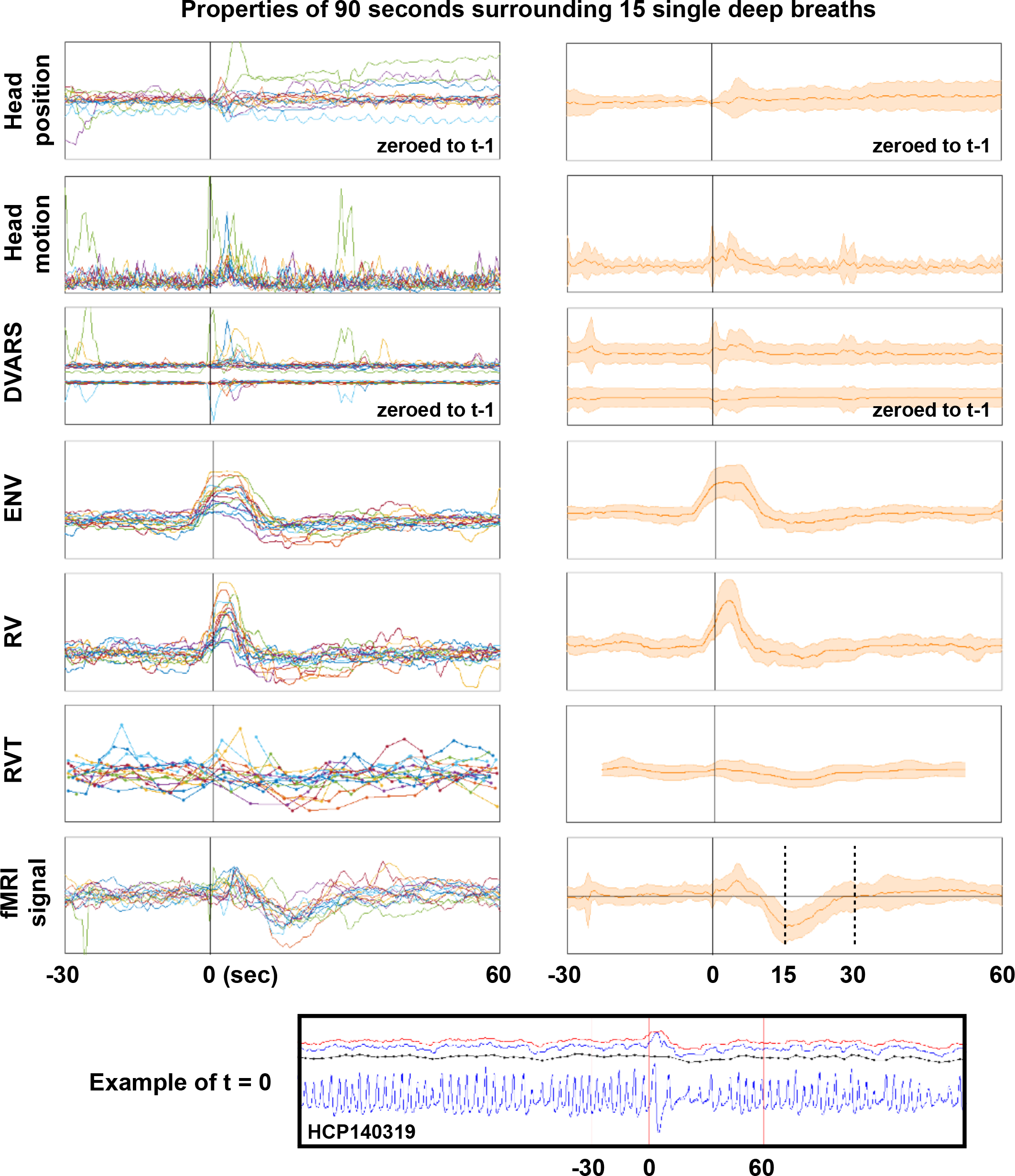
Summary of properties of 15 single deep breaths. Deep breaths were selected by respiratory traces and fMRI heat maps without knowledge of other properties, and full grayplots for all 15 scans are shown as a supplemental movie (these subjects are not in Figures 3 or 4). At bottom, an illustration of the respiratory trace and derived measures in a single subject, with vertical red lines indicating where t = 0 is set and the bounds 30 second prior to and 60 seconds after the breath. In the left column, data from each subject is shown in a different color, and traces for position, motion, DVARS, ENV, RV, RVT and mean gray matter signals (in minimally processed data) are plotted for 90 seconds surrounding deep breaths. In the right column, shaded plots show mean and standard deviations of the data. Axes are shifted slightly for the shaded head motion and DVARS plots to help visualization.

Thus, depending on the waveform and timing of a respiratory event, RVT may not mark an event, may mark it as a decrease, or may mark it as an increase. A corollary is that there is no universally satisfactory way to define peaks for RVT. Our standard settings will be to calculate RVT presuming respiratory cycles lasting 2 seconds or more. These settings may miss some rapid breaths, but will also avoid generating outliers in RVT of questionable accuracy. More generally – and this is an important point – there is no guarantee that a given respiratory event evident in the belt traces will be particularly noticeable in any of the derived respiratory measures. The measures are useful but imperfect indices of respiratory activity.

### Discordance of respiratory measures during single deep breaths

Single deep breaths are useful illustrative respiratory phenomena, for they occur in most subjects and have been studied as both instructed and spontaneous breaths (Birn et al., 2006; Power et al., 2018; Power et al., 2017b). All animals with lungs sigh routinely to counter the natural collapse of lung compartments (i.e., atelectasis), which is one reason deep breaths are common in scans (Li and Yackle, 2017). In resting state scans, deep breaths will probably usually represent sighs and yawns. To help convey the variety of respiratory waveforms denoting single deep breaths, 11 instances are shown in Figure 4 (there are thousands of examples in the online movies). In each of these instances, after an initial few seconds of moderate signal brightening, a black band in the gray plot reflects a pan-brain decrease in fMRI signals lasting until 20-30 seconds after the breath. There is almost always head motion associated with these breaths, often there is a DVARS spike in minimally preprocessed data, and there is sometimes, but sometimes not, a DVARS dip in FIX-ICA-denoised data. Note again that sometimes one or more respiratory measures do not capture the fact that something meaningful has occurred in the respiratory traces. Although ENV and RV often display bumps during the respiration, RVT sometimes displays bumps, sometimes displays nothing, and sometimes displays a dip, consistent with our previous discussion.

A different set of 15 deep breaths in 15 different subjects was selected by examining respiratory belt traces and grayscale fMRI signal heat maps (without knowledge of other properties); using only the traces and heatmaps let us to select isolated deep breaths that produced concomitant fMRI signal changes without actually knowing other properties of the scans. A gray plot for each of these 15 scans can be seen in an online movie^3^, with our markings of when the breaths began. For each breath, data from 30 seconds before and 60 seconds after the breath were extracted and are shown in Figure 5. Data for each of the different breaths are shown in a different color in the left column of Figure 5, and the right column summarizes the effects using mean and standard deviation shade plots. These events re-demonstrate the points above: 1) deep breaths cause ~30-second fMRI signal modulations, 2) deep breaths are marked by transient elevations of ENV and RV but usually not RVT, 3) deep breaths often exhibit motion, often exhibit step changes in head position, often exhibit DVARS spikes, and less often exhibit DVARS dips. The scales of the left and right columns are changes slightly to facilitate visualization.

Collectively, these observations demonstrate a disparity in the way various respiratory measures mark single deep breaths. The disparity arises chiefly because RVT is dependent on the timings and temporal characteristics of respiratory cycles: a larger-than-normal breath transpiring over a longer-than-normal time may appear quantitatively just like a typical breath occurring over a typical time. In contrast, the other measures are relatively insensitive to the exact temporal characteristics of the signal and are more sensitive to the amplitudes of the signal in the sampling window, which is why they routinely mark deep breaths.

As a final way to describe the tendencies of respiratory measure to identify deep breaths, the deep breaths of the first 10 and second 10 chronologically numbered subjects (total 20 hours of scanning) were visually identified, and each respiratory measure was scored as either clearly showing an abnormality (i.e., a bump or dip) or not clearly showing an abnormality at the times of clear deep breaths. In the first 10 subjects, in 21 clear deep breaths, ENV and RV marked 15 clearly (71%), and RVT marked 11 clearly (53%). In the next 10 subjects, of 48 clear deep breaths, 45 were marked clearly by ENV and RV (94%) and 24 by RVT (50%). These decisions underpinning these numbers are subjective but the statistics conform to the characteristics described above. The bottom line is that ENV and RV “miss” modest fractions of physiologically meaningful respiratory events, and RVT misses a substantially larger fraction of those same events.

### Concordance of respiratory measures during changes in breathing depth

Respiratory depth can have a major effect on all respiratory measures, and the effect is concordant. While maintaining regular, periodic breathing, subjects can breathe deeply, increasing exhalation of CO_2_, lowering pCO_2_, and thereby decreasing cerebral blood flow and causing BOLD signal decreases (black arrows of Figure 6). Periods of shallow breathing cause complementary phenomena (pink arrows of Figure 6). In the setting of respiratory depth modulation, high concordance is typically seen between ENV, RV, and RVT. The concordance follows from the sensitivity of all measures to changes in respiratory trace amplitude in the setting of relatively constant timings of cycles of breathing.

**Figure 6:**
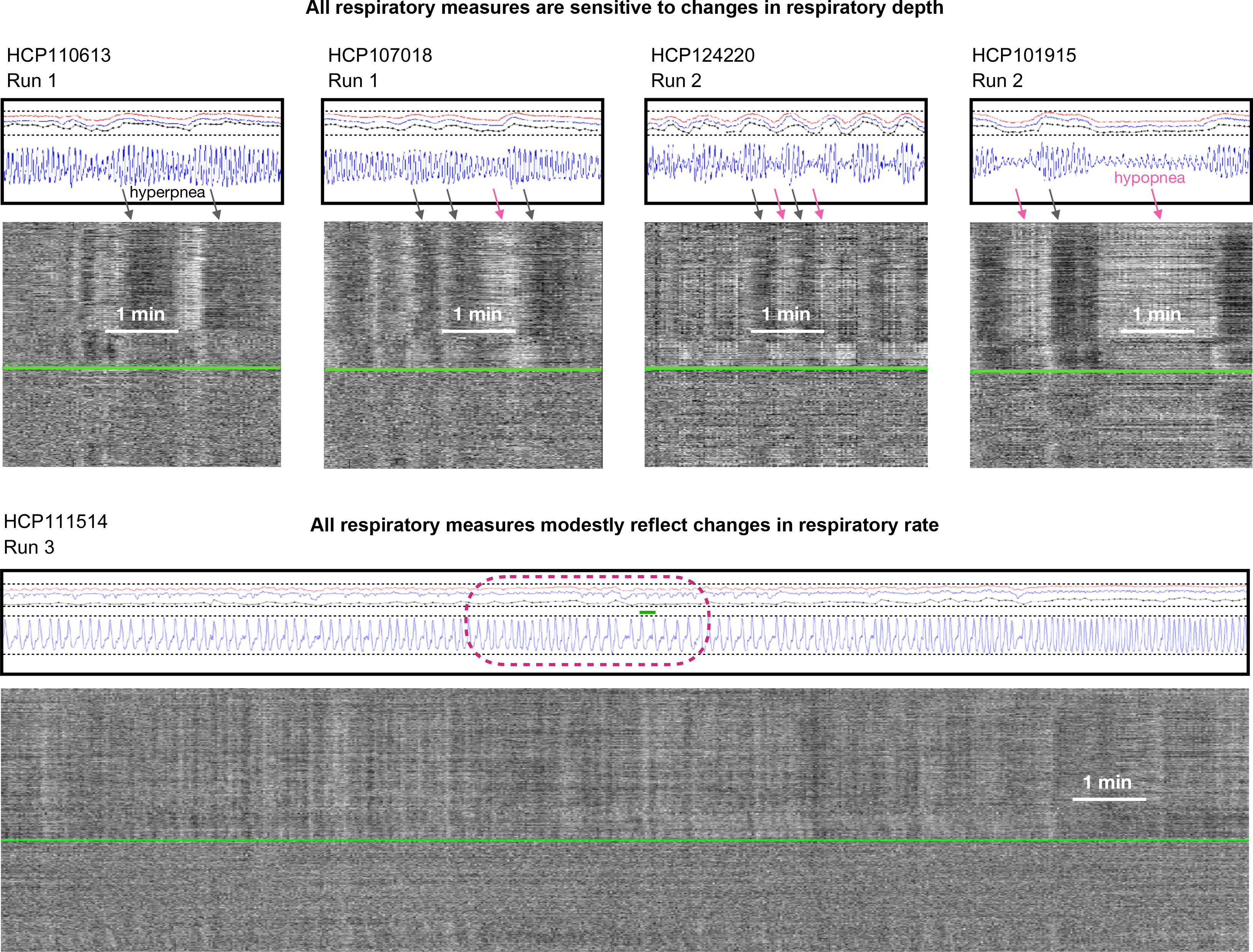
Examples of concordance between respiratory measures during sustained changes in respiratory depth and respiratory rate. At top, segments are shown from four subjects with substantial variation in respiratory depth, demonstrating large, concordant changes in respiratory measures in all scans. At bottom, a subject with much variation in respiratory rate (among the most variable we observed), with concordant but relatively modest modulation of respiratory measures by respiratory rate.

### Considerations about breathing rate

Breathing rate has a less obvious influence on respiratory measures than breathing depth, for one principal reason: although many subjects exhibit marked modulation of breathing depth, relatively few subjects show marked variation in respiratory rate within or across scans (Power et al., 2019a). The bottom row of Figure 6 illustrates one of the most variable subjects we encountered. There is an increase in all measures comparing the first few minutes to the final few minutes (compare via dotted lines), which one might attribute to respiratory rate increasing, but the amplitude has actually also increased (compare via dotted lines). The most informative comparison is seen in the middle of the scan (maroon dotted circle), where the respiratory rate falls almost by half while preserving amplitude: a decrease of respiratory measures with lowered respiratory rate is seen, but this decrease is much less than the decreases seen above with respiratory depth modulation.

Certain properties of the respiratory traces only become relevant at low respiratory rates. When occurring sufficiently rapidly, respiratory cycles are continuous oscillations (akin to sinusoidal forms), seen at the top of Figure 6 and in several other figures. However, when respiratory rates drop, this sinusoidal form does not hold, and there may be actual pauses in the respiratory cycle near the functional residual capacity (the resting point of the lung where recoil is balanced between expansion and contraction), rather than the continuous cycle of inspiration and expiration. This phenomenon is seen in the encircled portion of the bottom scan in Figure 6 where transient plateaus occur in the belt trace. When respiratory rates fall this low, distinctions among the respiratory measures emerge. Though RVT is sensitive to respiratory rates, precisely what happens in the waveform between the troughs and peaks is irrelevant (a sinusoidal cycle would have the exact same value as a flat line with a trough and a peak inserted at the appropriate times). In contrast, RV and ENV are windowed measures, and if the respiratory trace essentially pauses for some time, this pause can cause dramatic drops in the respiratory measure, depending on the relation of the pause to the windowing time. This sensitivity is seen in the RV trace in Figure 6, which exhibits marked dips during the pauses. Interestingly, the ENV measure is anticorrelated with RV at these times, displaying bumps during the plateaus. This bump is because of windowing – the ENV measures has 10 second windows (compared to 6 seconds for RV), and the respiratory rate is so low that at the midpoint between breaths the ENV window is encompassing one extra breath relative to when it is centered on a peak (small green line shows 10 second span). In contrast the RV window only ever encompasses a single cycle at respiratory rates this low.

The subject in Figure 6 was specifically chosen for demonstrating potential influences of respiratory rate. However, though differences in respiratory measures during especially slow breathing can be seen, this situation occurs infrequently, and respiratory rate modulation, when it occurs in HCP data, most often occurs at considerably higher frequencies (Power et al., 2019a), rendering the above considerations related to low rates largely academic, especially in comparison to the much more influential modulations of respiratory depth that are commonly seen in subjects.

### Systematic effects in respiratory measures over time

We turn now from properties of the respiratory measures in individual scans to properties at the group level. When ENV, RV, and RVT measures are examined over runs, a pattern emerges. Figure 7 shows the pattern for the RV measure (the same pattern is found in ENV and RV measures, see Figure S2). Nothing is easily seen in heatmaps of respiratory measures. But numerically, as shown in the red traces, the medians and means of respiratory measures across subjects decline over the course of each run, and the variance in those measures across subjects increases over the course of each run. To quantify these effects, values of these measures in minutes 1-4 and 11-14 were compared in each run by paired t-test (sampling once every 10 seconds to preserve independence in measures), yielding significant differences over time (green inset p values). To quantify the proportion of subjects displaying such effects, subjects were binned by the number of runs in which they displayed increases in the median, mean, and standard deviation within these time windows, and these proportions were fitted to binomial distributions, yielding a probability of 65% per run (95% C.I. 63%-67%) that measures would decrease and a 69% chance (95% C.I. 67%-71%) that variance would increase. The significance of this finding is that one should often expect systematic respiratory changes, and thus systematic BOLD respiratory effects, as fMRI runs progress.

**Figure 7:**
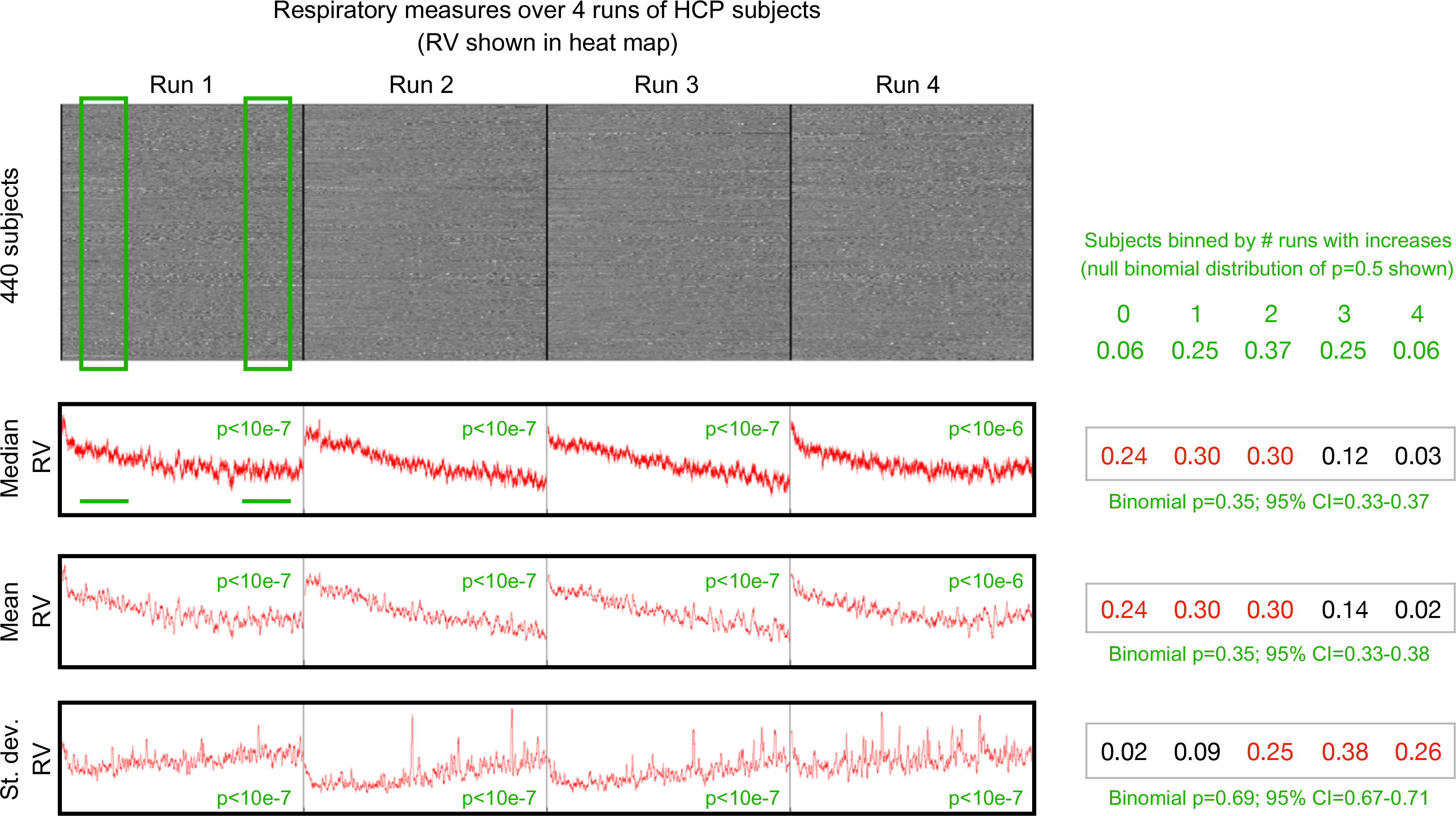
Respiratory measures systematically change during fMRI scans. At top, a heat map of the RV measures for all subjects with a complete set of 4 runs of fMRI data and acceptable physiologic measures. Median and mean RV values across subjects are shown, both showing decreases over time in each run. Standard deviation calculations across subjects show variability increasing over time in each run. To assess within-run changes, the mean, median, and standard deviation of RV was calculated in minutes 1-4 and 11-14 of each subject’s trace, in each run (see green boxes and underscores, samples were taken every 10 seconds). The green numbers by traces indicate the p values of paired t-tests of within-run comparisons of mean, median, and standard deviation in early and late time periods. At right, binning of subjects by the number of runs in which their within-run comparisons yielded increases is shown (raw number at top, fraction at bottom). A null binomial probability distribution is shown at top (for 50% chance of increase in a run), and each actual distribution is fit to a binomial distribution, with the probability and 95% confidence intervals shown in green. All intervals indicate an approximately 66% chance per run that a subject will show a decrease in RV and an increase in variance over each run. Similar findings are found using RVT and ENV measures (Figure S2).

Based on the properties noted above, decreases in breathing depth or rate could produce decreases in respiratory measure values over time. To test these possibilities, we calculated peak frequencies in minutes 1-4 and 11-14 of each run, and compared them within-subject by paired t-test. In each run, peak respiratory frequencies decreased in the later minutes compared to the earlier minutes, with small but significant effect sizes (p = 0.023, 3.0e-6, 0.002, and 6.6e-6 for the four runs). In the four runs, 18-25% of subjects showed an increase in rate, 39-46% showed a decrease in rate, and 34-42% showed no change in rate. We also examined the amplitude of the peaks that defined RVT within these early and late windows, and found that respiratory depth also decreased over each run (p = 1.4e-4, 3.4e-14, 4.6e-11, and 5.1e-5 for the four runs). In the four runs, 58-66% of subjects in each run exhibited decreases in amplitude. Thus, both reduced rate and reduced depth of breathing could each contribute to decreases in respiratory measures over time during these resting state scans, though depth likely has the greater influence.

## Discussion

In this paper we examined the characteristics of several respiratory measures used to index respiration and to flag respiratory events in fMRI studies. We used the HCP dataset to leverage its large number of subjects, the large amount of scan time per subject, and because this dataset has become a reference dataset for the field. Examination of hundreds of individual scans yielded three principal findings: 1) respiratory measures were broadly similar, especially RV and ENV, and all measures were prominently and similarly modulated by the depth of breathing; 2), there was a substantial discrepancy between respiratory measures when it comes to indexing deep breaths; and 3) respiratory measures change systematically across resting state scans. The principal implications of these finds are twofold and will be discussed further below: 1) RVT, by virtue of being insensitive to deep breaths, may fail to link relevant phenomena to respiration; 2) respiratory modulation of pCO_2_ ought to cause systematic BOLD changes over time in resting state scans.

### Implications of “missed” deep breaths

Deep breaths in resting state fMRI scans are probably mainly yawns and sighs. Sighs occur for a variety of reasons. “Physiological” sighs are universal among humans (and animals with lungs), they are estimated to occur about 12 times per hour in upright adults for lung-inflating purposes, they are even more frequent in the supine position, and they often include brief central apneas following the sigh (Li and Yackle, 2017). “Emotional” sighs (e.g., indicating disappointment) probably occur infrequently in resting state scans. Sighs can also be seen in the transition to sleep. In addition to sighs, the deep breaths in HCP resting state scans are also certain to also contain yawns, since subjects stared at a black screen for a quarter of an hour in a warm, dark environment. We know of no way to discriminate between these causes in the HCP data.

Deep breaths can be “missed” by any of the respiratory measures (Figure 3), but are more often “missed” by RVT than by RV or ENV (Figure 5). If deep breaths are not recognized by a respiratory measure, then a model using those measures will fail to assign “deep breath variance” to a respiratory cause and will instead assign it to anything modeled that co-occurs with deep breaths. Multiple physiological and mechanical changes accompany deep breaths, providing ample opportunity for such misattributions.

Deep breaths are often accompanied by head motion (or, relatedly, DVARS spikes or DVARS dips, see Figures 4 and 5). Deep breath fMRI signal modulations spanning ~30 seconds will be assigned to “motion” if either no respiratory cause is considered, or if the respiratory variable is insensitive to deep breaths. Such logic can explain why Byrge and colleagues (Byrge and Kennedy, 2018) find a 30-second “motion” signal following the larger motions of the HCP data with rather modest association to RVT. This logic could also explain why Glasser and colleagues (Glasser et al., 2018), even while attempting to index respiratory influences (via RVT), obtained associations of “motion” (DVARS dips) but not “respiration” (RVT) with ICA components whose spatial distribution resembles spatial maps of where deep breath respiratory effects are most prominent (Power, 2019).

A different version of the above scenario can occur with heart rate. Breathing is intimately related with heart rate via physical forces in the chest as well as by autonomic feedback loops. An example is the ongoing modulation of heart rate by breathing cycle seen in all humans, a phenomenon termed sinus arrhythmia, illustrated in the top row of Figure 8. With regard to deep breaths specifically, when subjects take deep breaths, it is common for the heart rate to increase transiently by 5-15 beats per minute before settling back to baseline or transient below-baseline values several seconds later. We documented this phenomenon previously in a different dataset (Power et al., 2017b), and routinely observe it in HCP scans. Figure 8 shows twelve individual examples heart rate increasing during and just after deep breaths, and also shows the effect in summary form for the set of deep breaths examined in Figure 5 (none of these breaths were chosen with knowledge of cardiac properties). By the same logic discussed above with motion, a study could assign deep breath fMRI signal changes to variables for heart rate or heart rate variability.

**Figure 8:**
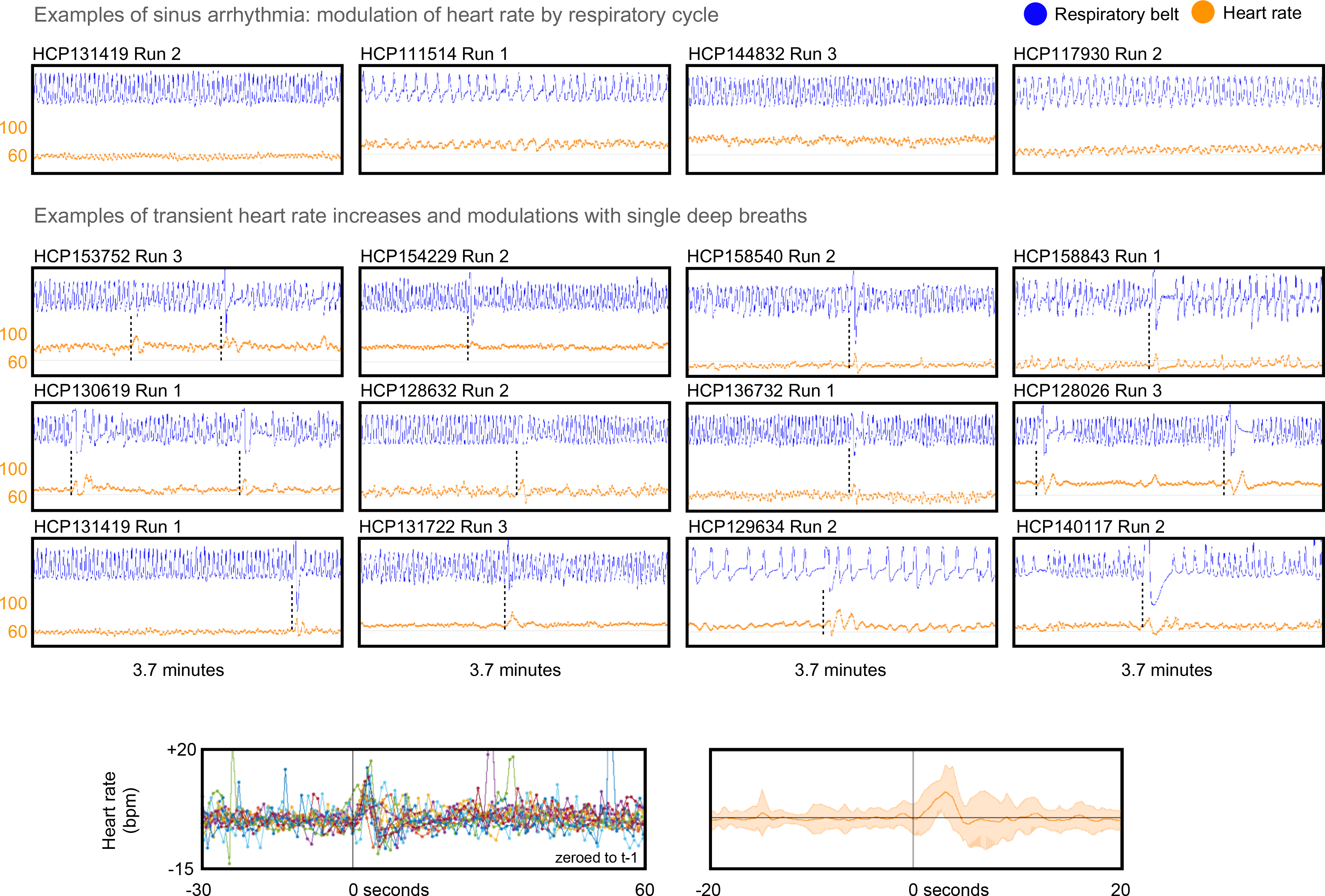
Examples of heart rate changes driven by respiration. At top, examples of sinus arrhythmia. Middle, examples of single deep breaths causing transient elevations and modulations of heart rate. At bottom, heart rate from the 90 seconds around 15 deep breaths examined in Figure 5 (only 13 subjects had usable cardiac records at the relevant time segment).

Such misassignment of respiratory variance can happen for any number of physiologic modulations that co-occur with deep breaths (e.g., pulse pressure). These scenarios are most likely to occur with RVT respiratory measures, but will occur even with ENV and RV to the extent they “miss” individual events.

### Implications of systematic changes in respiratory measures over time

Over the course of each 14.4-minute resting state scan of the HCP data, respiratory rates tended to decrease, respiratory depth tended to decrease, respiratory measures tended to decrease, and variance in respiratory measures tended to increase. Collectively, these observations indicate that investigators ought to expect pCO_2_ and thus BOLD signals to change systematically over a scan, at least at the group level.

Several reports in recent years have documented a “sensorimotor” set of ICA components (in spatial ICA in (Bijsterbosch et al., 2017) and temporal ICA in (Glasser et al., 2018)) with amplitudes that increase gradually across each of the resting state runs. As mentioned above, this “sensorimotor” distribution is the distribution in which respiratory models (e.g., of instructed deep breaths) explain the most variance (Birn et al., 2006; Birn et al., 2008; Wise et al., 2004). The ICA reports explained such effects as effects of arousal, which may be true, but the most direct explanation of the component properties would be as consequences of respiratory phenomena in the scans. Stated differently, these components may be signatures of arousal only insofar as states of arousal cause or correlate with altered respiratory properties (Power, 2019; Power et al., 2017a).

It is of some interest that a large study that monitored sleep state during resting state fMRI scanning reported that approximately two thirds of subjects showed evidence of sleep or decreased arousal as a scan progressed (Tagliazucchi and Laufs, 2014). Two thirds is also the fraction of the HCP subjects examined here who exhibited decreased respiratory measures and increased respiratory variance over their scans. Studies in typical human adults document that the transition from wakefulness to non-rapid-eye-movement sleep is characterized by a mild reduction (~10%) in respiratory rate and a mild reduction in tidal volume (~5%) (Berssenbrugge et al., 1983), which would accord with our reduced rates and amplitudes over time. A related set of observations is that several studies of sleep have also recovered a “sensorimotor” set of signal changes that denote changes in arousal and sleep state (Horovitz et al., 2008; Tagliazucchi and Laufs, 2014; Tagliazucchi et al., 2013). One might speculate that these fMRI signatures of sleep may in large part actually be respiratory signals, by the same logic mentioned in the previous paragraph. One could perhaps make a counterargument that the modeling of spontaneous respiration is itself actually modeling arousal, but such an argument is undermined by the fact that instructed and uninstructed breaths produce similar temporal and spatial patterns in fMRI signals (Birn et al., 2008; Wise et al., 2004).

### On monitoring respiration

It is ironic that the one fMRI respiratory measure (RVT) that attempts to incorporate proxies of standard physiologic respiratory measures (e.g., inspired volume, cycle time) performs relatively poorly compared to measures that are simply ad hoc descriptions of a waveform (RV and ENV). This poorer performance is partially explained by practical considerations about respiratory belt records. RV and ENV are relatively insensitive to the exact timings of wave modulations by virtue of their windowed calculation, and so brief non-respiratory motions or other artifactual disruptions of the belt record are less problematic for those measures than for RVT, which strongly depends on times between peaks in the trace. But even absent such factors, the parameter indexed by RVT – air moved in a cycle time – is by definition relatively insensitive to outlier breath volumes so long as they scale with breath times. We experimented with a number of other “physiologically motivated” computations based on the belt records, but failed to obtain a parameter sufficiently reliable to bother reporting. Our view is that of the present parameters, ENV and RV are similar in practice, are easy and unambiguous to compute, and usually “flag” noticeable events in the respiratory timeseries. Clearly there is room for improvement in these measures (given that they can “miss” deep breaths), but it seems that a substantial limiting factor in deriving quality respiratory measures is a lack of reliable and comprehensive records to begin with. For these and other reasons, we have begun to collect a longitudinal multi-echo, multi-band fMRI dataset with comprehensive physiologic recording that will eventually be made available to the public, including simultaneous capnography, inductance plethysmography (thus including an abdominal belt for comparability to “single abdominal belts”), electrocardiography, pulse oximetry with gas saturations, galvanic skin responses, and pupillometry and eye tracking. With such data in hand, one hopes there would be little obstacle to creating precise respiratory models, at least from a reliability or comprehensiveness standpoint.

### Conclusions

Respiration can cause a variety of effects in fMRI data, from alterations of cerebral blood flow and BOLD signal to real head motions to pseudomotion. Breathing is also yoked to cardiovascular parameters via the physics of the chest and by autonomic loops. There are thus many reasons an investigator might wish to monitor respiration during fMRI scans. However, making use of such respiratory records is not trivial, and the different respiratory measures that can be derived from the records can perform quite differently at indexing respiration and respiratory events depending on the kinds of breathing in a scan. We studied task-free scans of young adults here; similar analyses in other age ranges and in task settings may be of interest. The field would benefit from more robust and reliable measurement of respiration, both in terms of physiological monitoring at the scanner and in terms of deriving variables from the physiological records.

## Acknowledgments

This work was supported by Simons Foundation Grant 528440 and a gift from the Mortimer D. Sackler, M.D. family. We thank Kevin Tran and the NIH Helix/Biowulf staff for computing support.

## SUPPLEMENTAL

**Figure S1:**
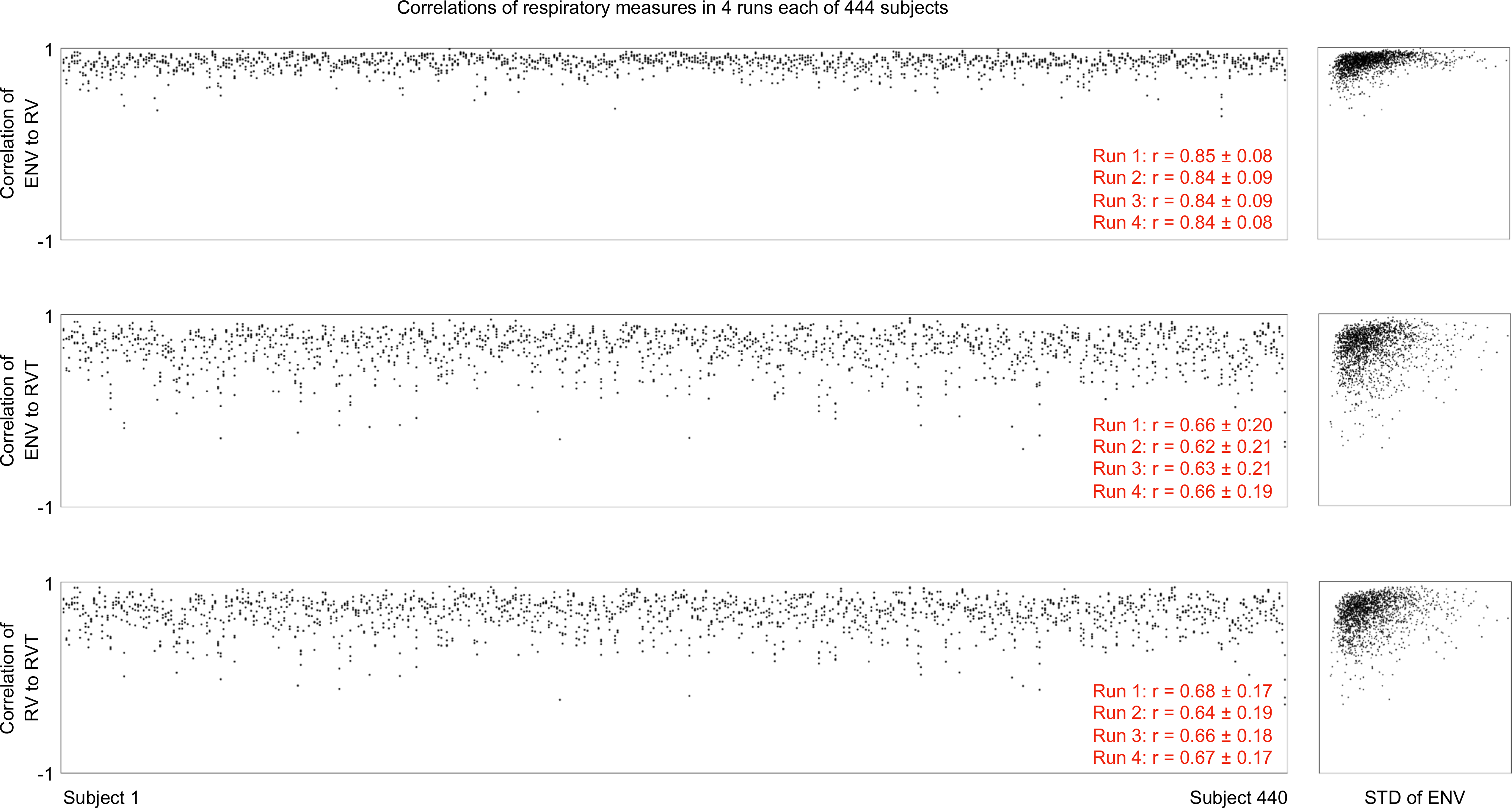
Correlations among the 3 respiratory measures for each run of each subject, and summary measures for each run. At right, all correlations are plotted as a function of the standard deviation of the ENV trace.

**Figure S2:**
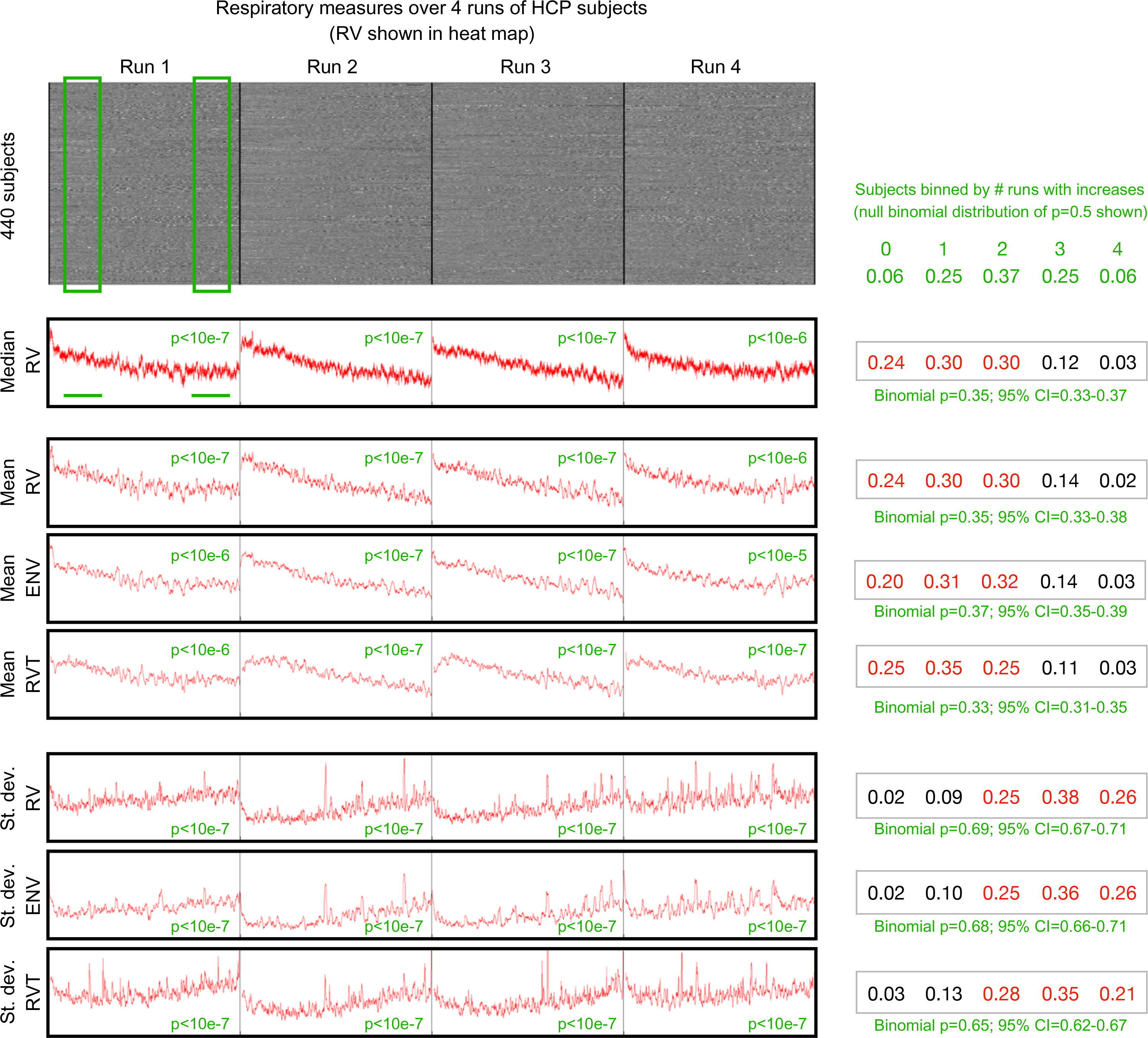
Across-run effects in respiratory measures. At top, a heat map of the RV measures for all subjects with a complete set of 4 runs of fMRI data and acceptable physiologic measures. Median and mean RV values across subjects are shown, and have the same form, indicating that mean values are representative summary measures. Mean RV, ENV, and RVT values are then shown across subjects, all showing decreases over time in each run. Standard deviation calculations across subjects show variability increasing over time in each run. To assess within-run changes, the mean, median, and standard deviation of each measure (RV, ENV, and RVT) was calculated in minutes 1-4 and 11-14 of each subject’s trace, in each run (see green box and underscores, samples taken every 10 seconds). The green numbers by traces indicate the p values of paired t-tests of within-run comparisons of mean, median, and standard deviation in early and late time periods. At right, binning of subjects by the number of runs in which their within-run comparisons yielded increases is shown (raw number at top, fraction at bottom). A null binomial probability distribution is shown at top (for 50% chance of increase in a run), and each actual distribution is fit to a binomial distribution, with the probability and 95% confidence interval shown in green. All intervals indicate an approximately 66% chance per run that a subject will show a decrease in ENV/RV/RVT and an increase in variance in those measures.

## Footnotes

1 www.jonathanpower.net/2018-glasser-comment.html

2 www.jonathanpower.net/2019-respiratory-measures.html

3 www.jonathanpower.net/2019-respiratory-measures.html

